# Endosomal Trafficking of Two Pore K^+^ Efflux Channel TWIK2 to Plasmalemma Mediates NLRP3 Inflammasome Activation and Inflammatory Injury

**DOI:** 10.1101/2022.10.12.511914

**Authors:** Anke Di, Long Shuang Huang, Bisheng Zhou, Peter T. Toth, Yamuna Krishnan, Asrar B. Malik

## Abstract

Potassium efflux via the two pore K^+^ channel TWIK2 is a requisite step for the activation of the NLRP3 inflammasome, however it is unclear how the efflux is activated in response to cues. Here we report that during homeostasis, TWIK2 resides in endosomal compartments. TWIK2 is transported by endosomal fusion to the plasmalemma in response to increased extracellular ATP resulting in extrusion of K^+^ ATP-induced endosomal TWIK2 plasmalemma translocation is regulated by Rab11a. Deleting Rab11a or ATP ligated purinergic receptor P2X7 prevented endosomal fusion with the plasmalemma and K^+^ efflux and NLRP3 inflammasome activation in macrophages. Adoptive transfer of Rab11a-deleted macrophages into mouse lungs prevented NLRP3 inflammasome activation and inflammatory lung injury. Rab11a-mediated endosomal trafficking in macrophages thus regulates TWIK2 abundance and activity on the cell surface and downstream activation of the NLRP3 inflammasome. Endosomal trafficking of TWIK2 to the plasmalemma is therefore a potential therapy target in acute or chronic inflammatory states.

## Introduction

Inflammasomes are key components of the immune system and inflammatory signaling platforms responsible for detecting injury mediators released during infection and tissue damage and thereby initiate the inflammatory response^1–4^ NLRP3 (<u>N</u>ucleotide-binding oligomerization domain-<u>L</u>ike <u>R</u>eceptor containing <u>P</u>yrin domain <u>3</u>) inflammasome expressed in immune cells such as macrophages is a key determinant of acute immune responses such as acute lung injury and COVID-19^4–6^ as well as chronic inflammatory diseases such as atherosclerosis, cancer or metabolic syndrome^3,7^. Activation of NLRP3 inflammasome complex is a multi-step process involving assembly of key proteins, activation of caspase-1 which cleaves pro-Interleukin-1β to release the active form of this inflammatory cytokine^5,8^. NLRP3 inflammasome consists of a sensor (NLRP3), an adaptor (apoptosis-associated speck-like protein containing a caspase recruitment domain - ASC), and an effector (caspase 1)^1–4^. Oligomerized NLRP3 recruits ASC which in turn recruits caspase 1 and enables proximity-induced caspase 1 self-cleavage and activation^1–4^ Several studies have elucidated the structure and assembly mechanisms of the NLRP3 complex. The inactive NLRP3 form (double-ring cages of NLRP3) is predominantly membrane associated (such as endoplasmic reticulum - ER, mitochondria and Golgi Apparatus) and is recruited, assembled and activated at the centrosome^9–17^ However, little is known about the early triggers initiating the assembly and activation of NLRP3 complex.

An essential mechanism of NLRP3 assembly is the efflux of potassium (K^+^) at the plasmalemma^18–20^ through the potassium channel TWIK2 (the <u>Tw</u>o-pore domain <u>W</u>eak <u>I</u>nwardly rectifying <u>K^+^</u> channel 2), a member of the two-pore domain K^+^ channel (K_2P_) family (K_2P_ 6.1, encoded by *Kcnk6*)^21,22^ Efflux of potassium generates regions of low intracellular K^+^ which promote a conformational change of inactive NLRP3 to facilitate NLRP3 assembly and activation^23^. TWIK2 mediated plasmalemmal potassium efflux thus serves as a checkpoint for the initiation of adaptive host defense as well as maladaptive inflammatory signaling mediated by NLRP3^22^ This essential function of TWIK2 mediated K^+^ efflux in initiating macrophage inflammatory response raises the question how TWIK2 activity is fine-tuned to avoid premature or intracellular triggering of NLRP3 during homeostasis while at the same time providing for rapid TWIK2 activation mechanism in response to extracellular tissue damage. Due to the high gradient of K^+^ across the plasmalemma, the presence of TWIK2 at the plasma membrane may result in basal potassium efflux even and thus lead to inappropriate inflammasome activation. Therefore the question of fine control of K^+^ efflux becomes even more important.

It has been shown that the activity of several ion channels is regulated by endosomal trafficking from the cytosol to the plasmalemma such as the G protein-activated inwardly rectifying K^+^ (GIRK) channels^24–26^ and cardiac pacemaker channels - hyperpolarization-activated cyclic nucleotide-gated (HCN) ion channels HCN2 and HCN4^27^. Here we investigated whether TWIK2 is similarly sequestered in the steady-state in cytosolic endosomal compartments to shield cells from maladaptive inflammasome activation and only trafficked to the plasmalemma during cues elicited by tissue injury to trigger potassium efflux and activate NLRP3.

Using electron microscopy and electrophysiological studies, we found that TWIK2 K^+^ channel in macrophages was expressed in endosomes at rest and was translocated to the plasmalemma upon extracellular ATP challenge, an indicator of tissue damage. The Ca^2+^ sensitive GTP binding-protein Rab11a was responsible for endosomal TWIK2 translocation to the plasmalemma. Furthermore, Inhibition of endosomal fusion with the plasma membrane prevented NLRP3 inflammasome activation. We also showed that adoptive transfer of Rab11-deleted macrophages into mouse lungs prevented NLRP3 inflammasome activation and inflammatory lung injury. Our studies thus demonstrate an essential mechanism by which endosomal trafficking of TWIK2 and K^+^ efflux trigger NLRP3 inflammasome activation. The results point to inhibition of endosomal plasmalemma fusion as a potential anti-inflammatory therapy target.

## RESULTS

### Endosomal TWIK2 plasmalemmal translocation post ATP challenge in macrophages

TWIK2 belongs to the constitutively active K2P background potassium channel family^21^; however, TWIK2 plasmalemmal current in macrophages is only observed following challenge with extracellular ATP^22^ A possible explanation is that TWIK2 is not present at the plasmalemma during homeostasis basal state and is only trafficked to the plasmalemma in response to selective environmental cues, similar to what has been reported for other ion channels activity^24–27^ To test this question, we visualized the intracellular TWIK2 plasmalemma translocation upon ATP challenge by expressing TWIK2-GFP in macrophages. TWIK2-GFP plasmids were transfected into RAW 264.7 macrophages for 48h and the cells were imaged with confocal microscopy before and after challenge with extracellular ATP to mimic tissue injury. TWIK2 translocated towards the plasmalemma within 2 min after extracellular ATP addition (**Fig. 1A, Supplemental video1**). To assess TWIK2 plasmalemma insertion after ATP challenge, we examined the cellular distribution of TWIK2 before and after ATP challenge using confocal microscopy and immunogold labelled electron microscopy. Confocal images showed TWIK2 intracellular distribution before the ATP challenge, and clear TWIK2 plasmalemmal translocation after ATP challenge (**Fig. 1B**). Plasmalemmal translocation of TWIK2 was confirmed by immunogold-labeled electron microscopy (**Fig. 1C, Supplemental Fig. 1**). Intracellular TWIK2 plasmalemmal translocation was ATP concentration-dependent (**Fig. 1D**). To identify the location of intracellular TWIK2, we labeled TWIK2 with fluorescent TWIK2 antibody combined with fluorescence labeled markers for early endosomes (EE) with EEA1 (Early Endosome Antigen-1) antibody, recycling endosomes (RE) with Rab11a antibody, lysosomes with LAMP1 (Lysosomal-Associated Membrane Protein 1) antibody, and endoplasmic reticulum (ER) with PDI (Protein Disulfide Isomerase) antibody. TWIK2 was only present in endosomes (both EE and RE) whereas lysosomes or ER were free of TWIK2 (**Fig. 2**). Thus, TWIK2 is primarily distributed in endosomes during homeostasis and is translocated to the plasmalemmal upon ATP challenge.

**Figure 1.**
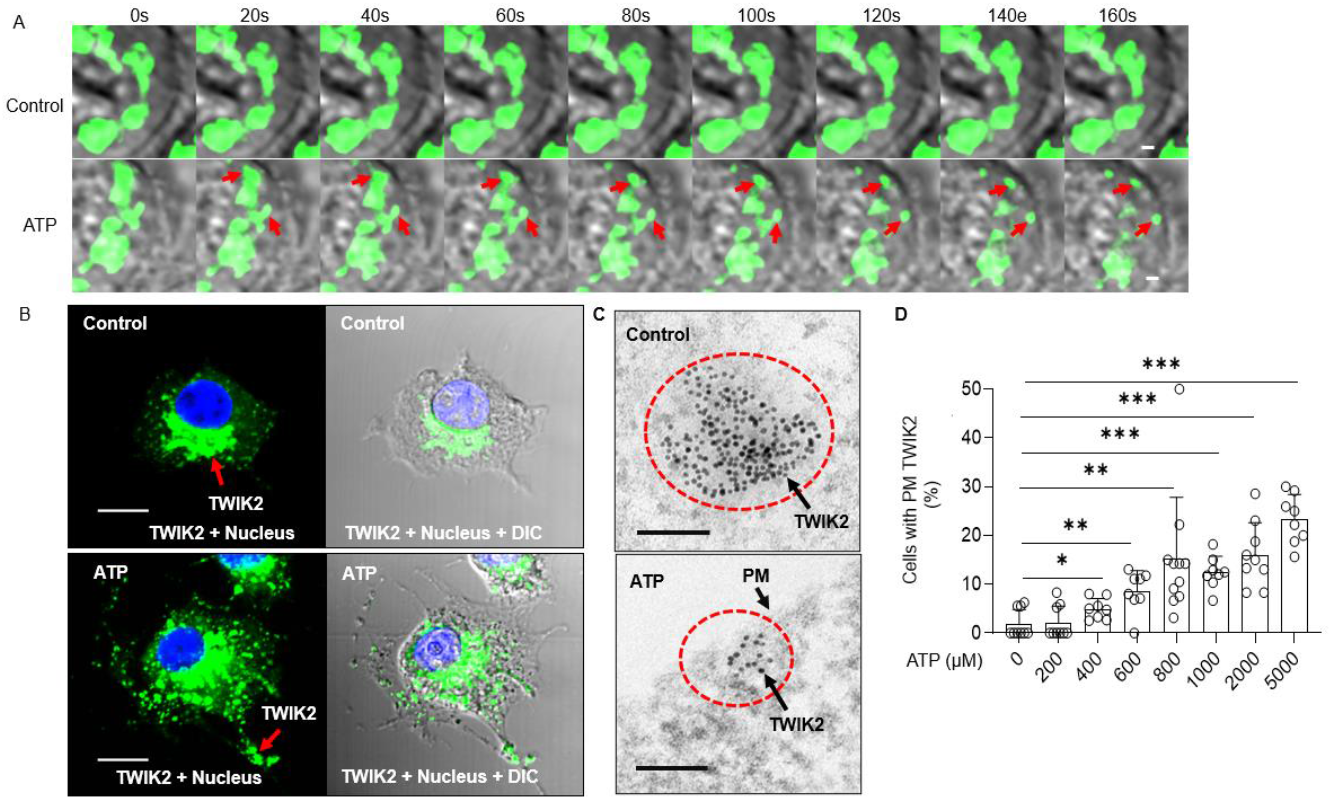
Intracellular TWIK2 plasmalemma translocation upon ATP challenge. **A.** Visualization of intracellular TWIK2 plasmalemma translocation upon ATP challenge. Representative confocal images of macrophages with TWIK2-GFP from 3 independent experiments. Representative videos of TWIK2-GFP plasmalemma translocation upon ATP challenge in these experiments were shown in ***Supplemental videos*.** TWIK2-GFP plasmid was transfected into RAW 264.7 cells for 48hr and cells were imaged with confocal microscope in the presence or absence of extracellular ATP (5mM). Red arrows showing the movement of TWIK2 (green) toward plasma membrane. Scale bar = 1μm. **B.** Representative confocal images of TWIK2 immunostaining from 3 independent experiments in the mouse RAW 264.7 macrophage cell. TWIK2 distribution before (upper panel) and after (lower panel) ATP (5 mM, 30min) challenge was determined with fluorescent immunostaining with anti-human TWIK2 (aa71-120) antibody (Life Span Bioscience, LSBio #LS-C110195-100) imaged with confocal microscope. Scale bar = 10μm. **C.** Confirmation of TWIK2 plasma membrane translocation from immunogold labeling electron microscope before (upper panel) and after ATP (lower panel) (5 mM, 30min) challenge of RAW 264.7 macrophages. TWIK2 (10 nm gold particles) was identified with anti-TWIK2 antibody as in **B**. Scale bar = 100nm. Note the vesicular structure outlined by immunogold in the upper panel and distribution of immunogold labeled TWIK2 in the plasma membrane (PM) after ATP challenge in the lower panel. See more details in ***Supplemental Figure 1*. D**. ATP concentration-dependent TWIK2 plasma membrane translocation in mouse monocyte derived macrophages (MDMs). TWIK2 plasma membrane (PM) translocation was analyzed based on confocal images from experiments performed as shown in **B**. \**P*<0.05, \*\**P*<0.01, \*\*\**P*<0.001 compared with control (0 ATP) group, n = 8).

**Figure 2.**
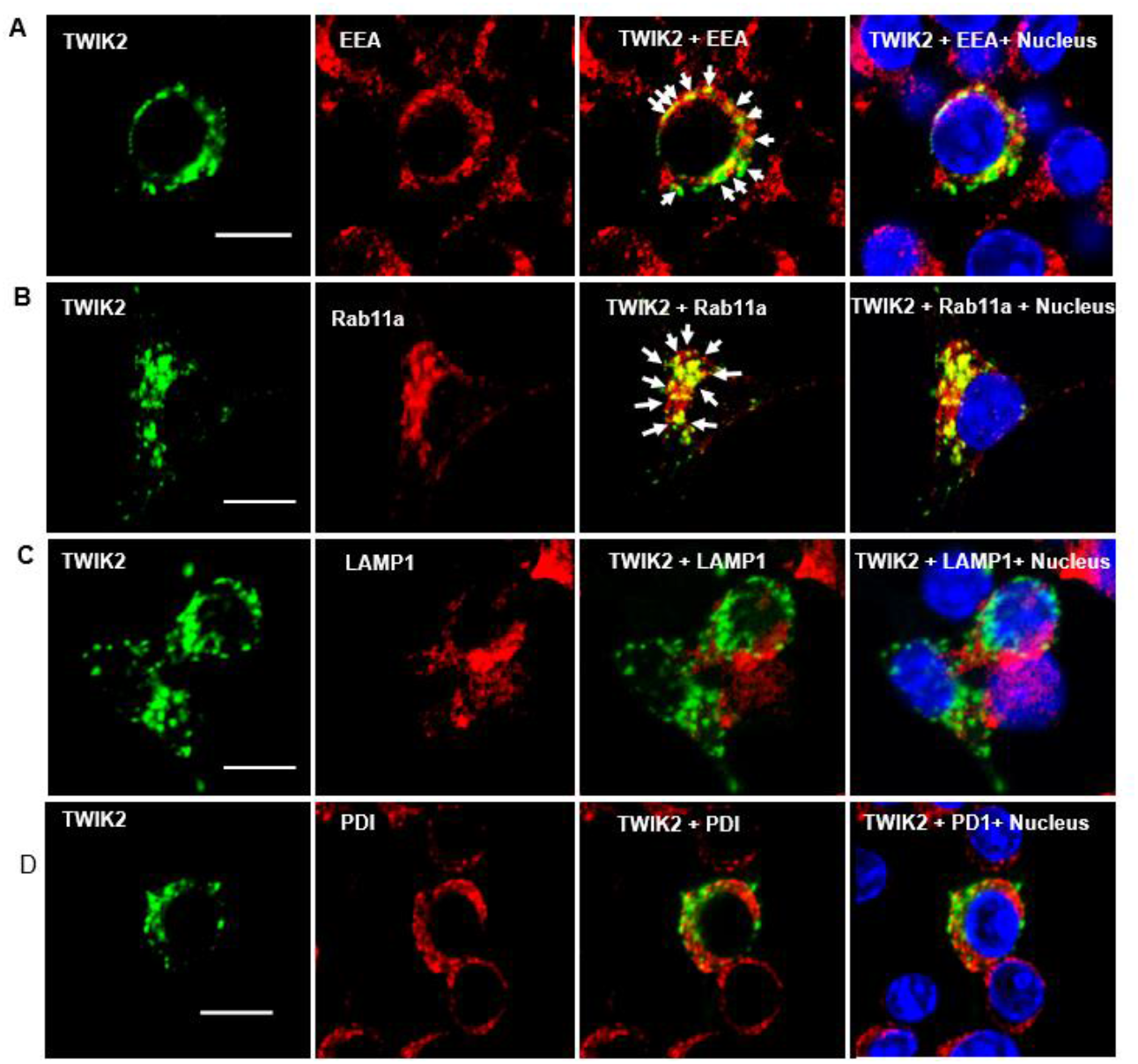
Endosomal localization of intracellular TWIK2 determined with fluorescent immunostaining of macrophages. Representative confocal images of RAW 264.7 macrophages with TWIK2-GFP from 3 independent experiments. TWIK2 intracellular localization was determined with fluorescent immunostaining with TWIK2 antibody along with other various antibodies against some specific vesicular proteins and imaged with confocal microscope. RAW 264.7 macrophages were fixed/permeabilized followed by immunostaining. TWIK2 (green) was identified with specific TWIK2 antibody (LSBio #LS- C110195-100) in **A** through **D**. Early Endosomes (EE, red) were identified with specific antibody (clone1D4B) against EEA1 (C45B10 from Cell Signaling Technology) in **A**. Recycling Endosomes (RE, red) were identified with antibody against Rab11a (ab65200 from Abcam) in **B**. Lysosomes (red) were identified with antibody against lysosomal membrane protein LAMP1 (D2D11 from Cell Signaling Technology) in **C**. ER was identified with antibody against PDI (C81H6 from Cell Signaling Technology) in **D**. Note the localization of TWIK2 (green) in both the EE and RE (white arrows) but not in lysosomes or ER. Scale bar = 10μm.

### ATP-induced exocytosis induces plasmalemma potassium efflux and NLRP3 inflammasome activation in P2X7- and Ca^2+^- dependent manner

Endosomal distribution of TWIK2 and plasmalemmal translocation suggest a mechanism of plasmalemma insertion of TWIK2 upon ATP challenge. To address TWIK2 distribution we monitored exocytosis (the fusion event of an intracellular vesicle with plasmalemma) through membrane capacitance measurements reflecting increased membrane surface area upon fusion of vesicles with the plasmalemma^28,29^ (**Fig. 3A**). We observed ATP induced exocytosis as reflected by the increase of plasmalemma capacitance measurement in monocyte-derived macrophages (MDMs) (**Fig. 3A**). ATP-induced exocytosis determined by capacitance increase was inhibited by either deletion of P2X7 (*P2×7^−/−^*), absence of extracellular Ca^2+^ or by using the vesicle plasma fusion inhibitor (Vacuolin)^30,31^ in MDMs (**Fig. 3A**).

**Figure 3.**
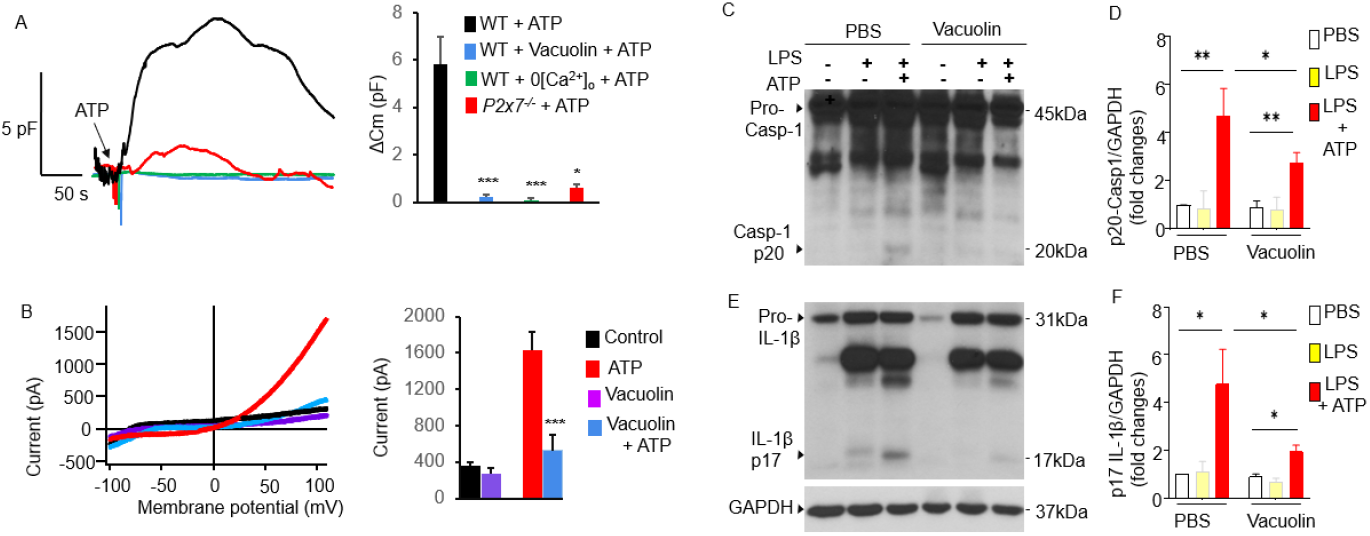
P2X7 dependent ATP-induced exocytic event is linked to plasmalemma potassium efflux and NLRP3 inflammasome activation. **A-B.** Exocytic event is linked to plasmalemma potassium efflux. **A**. Exocytic event was evaluated with measurement of whole cell plasma membrane capacitance (Cm) which reflects the membrane surface changesan. Left panel: Raw Cm traces recorded in monocyte derived macrophages (MDMs) from either WT or *P2×7^−/−^* mice under different conditions: 0 extracellular Ca^2+^ (0[Ca^2+^]_o_) to confirm the involvement of P2X7 which mediated Ca^2+^ influx; Vacuolin, an inhibitor of vesicle plasma membrane fusion to confirm the involvement of membrane fusion events. 5 mM ATP was added as indicated. In the Vacuolin group, the cells were treated with 10 μM Vacuolin for 2 hrs before challenged with ATP. ATP treatment causes Cm increase, indicating intracellular vesicle fusing with plasma membrane. Right panel: summary of capacitance changes shown in the left panel. *p < 0.05, ***p < 0.001 compared with WT + ATP group, n = 3). Note: vesicle membrane fusion inhibitor Vacuolin inhibits ATP-induced exocytosis (Cm increase), indicating the increased Cm caused by ATP challenge results from the fusion of intracellular vesicles with the plasma membrane. Also note that the fusion event is P2X7 and extracellular Ca^2+^ dependent. **B**. Vesicle-plasmalemma fusion dependent of ATP-induced potassium efflux current. Whole cell current was recorded with patch clamp in MDM with or without vacuolin (10 μM). Currents were elicited with a ramp voltages running from −110 mV to +110 mV within 200 ms applied to cells with an interval of 1 s. Cells was holding at 0 mV. Cells were bathed in solutions with K^+^ as the major outward current and Na^+^ and Ca^2+^ as the major inward current. Left panel: Representative I-V plot of whole cell current in MDM. Right panel: Summary from experiments displayed in the left panel. ***p < 0.001 compared with ATP group, n = 6). Note: Cells pretreated with inhibitor of vescular fusion protein Vacuolin showed significantly decreased current induced by ATP. **C-F**. Inhibition of vesicle plasmalemma fusion prevents NLRP3 inflammasome activation in macrophages. **C**, **E**: Representative results of Western blot from three independent experiments showing reduced Caspase 1 activation (reduced Casp-1 p20, **C**) and IL-1β maturation (reduced IL-1β p17, **E**). MDMs pretreated with vesicle plasmalemma fusion inhibitor vacuolin (10 μM, 2hr) were primed with LPS (3 hr) and subsequently challenged with ATP (5 mM) for 30 min. Cell lysates or pellets were immunoblotted with indicated antibodies (anti-TWIK2 or anti-IL1β). **D**, **F**: Quantification of results shown in **C**, **E**. *p < 0.05 **p < 0.01, n = 3. Note the reduced Casp-1 p20) and IL-1β p17 cells treated with vacuolin.

To address the role of vesicle plasmalemma fusion and Ca^2+^ in TWIK2 activation and NLRP3 inflammasome activation, we next investigated whether reducing extracellular Ca^2+^ or inhibiting ATP-induced vesicle-plasmalemma fusion altered ATP-induced potassium current and NLRP3 inflammasome activation. We demonstratef the requisite role for extracellular Ca^2+^ and plasmalemma-endosome fusion in ATP-induced NLRP3 inflammasome activation. In these experiments, caspase 1 activation (indicated by p20 derived from pro-caspase 1) and IL-1β maturation (indicated by p17 derived from pro-IL-1β) were used as the indicator of NLRP3 inflammasome activation. First, we tested the effects of Vacuolin and BAPTA (1,2-Bis [2-aminophenoxy] ethane-N,N,N’,N’-tetraacetic acid tetrakis [acetoxymethyl ester]) in the ATP-induced plasmalemma K^+^ current and NLRP3 inflammasome activation. Cells pretreated with Vacuolin or BAPTA showed a significant reduction in ATP-induced current (**Fig. 3B** and **Fig. 4AB**) and NLRP3 inflammasome activation (**Fig. 3C-F and Fig. 4C**). Second, we examined the role of Ca^2+^ in ATP-induced NLRP3 inflammasome activation by reducing extracellular Ca^2+^. Both caspase 1 activation and IL-1β maturation were significantly reduced in MDMs treated with ATP (5 mM) in the absence of extracellular Ca^2+^ (**Fig. 4D-G**). These results together showed a key role of P2X7-mediated Ca^2+^ influx in ATP-induced exocytosis linked to plasmalemma potassium efflux and NLRP3 inflammasome activation.

**Figure 4.**
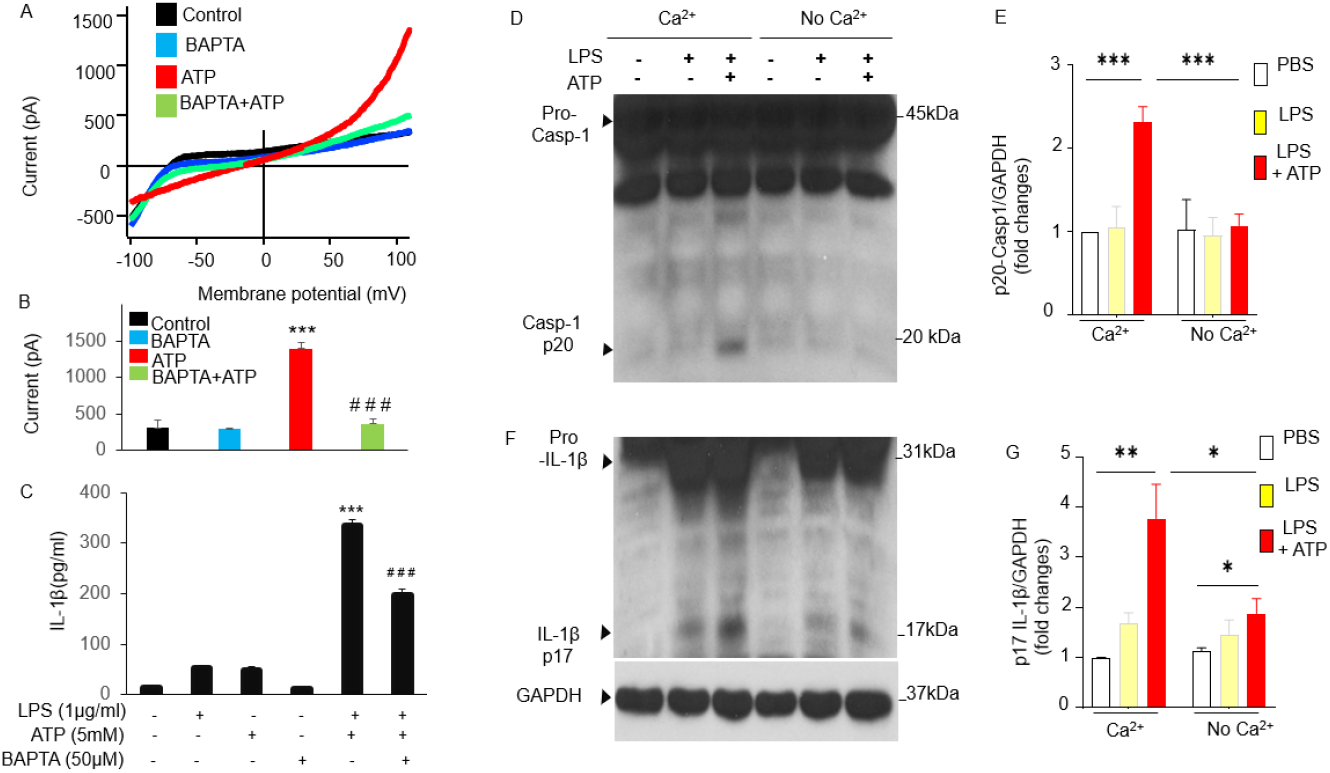
ATP-induced potassium current and NLRP3 inflammasome activation in Cam-dependent manner. **A-B**. Ca^2+^ dependent of ATP-induced potassium efflux current. Whole cell current was recorded with patch clamp in MDM with or without pretreatment of BAPTA-AM (1,2-Bis [2-aminophenoxy] ethane-N,N,N’,N’-tetraacetic acid tetrakis [acetoxymethyl ester], 10 μM) for 20min. Currents were elicited with a ramp voltages running from −110 mV to +110 mV within 200 ms applied to cells with an interval of 1 s. Cells was holding at 0 mV. Cells were bathed in solutions with K^+^ as the major outward current and Na^+^ and Ca^2+^ as the major inward current. **A**. Representative I-V plot of whole cell current in MDM. **B**. Summary from experiments displayed in **A**. ***p < 0.001 compared with control group, n = 7;^###^p < 0.001 compared with ATP group, n = 7). Note: Cells pretreated with BAPTA-AM showed significantly decreased current induced by ATP (5mM). **C**. Reduced IL-1β release in the presence of Ca^2+^ chelator BAPTA-AM in the supernatants measured using ELISA method (***p < 0.01 compared with LPS or ATP group;^###^p < 0.001 compared with LPS+ATP group, n = 3). Note: Cells pretreated with BAPTA-AM showed significantly decreased current induced by both ATP (5mM). **D-G**. Extracellular Ca^2+^dependent NLRP3 inflammasome activation in macrophages. MDMs were primed with LPS and subsequently challenged with ATP and cell lysates or pellets were immunoblotted with indicated antibodies (anti-Caspase 1 or anti-IL1β). **D**, **F**: Representative Western blotting results from three independent experiments showing reduced Caspase 1 activation (reduced Casp-1 p20) and IL-1β maturation (reduced IL-1β p17) in the absence of extracellular Ca^2+^. **E**, **G**: Quantification of results shown in **D**, **F**. *p < 0.05 **p < 0.01, ***p < 0.001, n = 3. The absence of extracellular Ca^2+^ prevented ATP-induced NLRP3 inflammasome activation in MDMs.

### Rab11a in recycling endosomes is required for endosomal TWIK2 plasmalemmal translocation, sepsis-induced NLRP3 inflammasome activation, and lung inflammation

Since plasmalemmal translocation of endosomal TWIK2 involves endosomal fusion with the plasmalemma as described above, we focused on identifying the Ca^2+^ sensitive GTP-binding protein translocation and fusion machinery (Rab family)^32,33,34^, Synaptotagmin family (Syt)^35^, and Vesicle associated membrane proteins (Vamp, a.k.a. Synaptobrevin)^36–38^. Quantitative assessment of expression showed that mRNA levels of Rab11a were the highest as compared to other genes involved in the translocation and fusion of endosomes (**Supplemental Fig. 3**). Since Rab11 a is generally thought to regulate the function of specific endosomal subpopulation, the recycling endosomes^39,40^, we examined the role of Rab11a in mediating endosomal TWIK2 plasmalemma translocation. We first determined the cellular distribution of Rab11a before and after ATP challenge using fluorescence immunostaining confocal microscopy. Images showed Rab11a plasmalemmal translocation after ATP challenge in macrophages (**Fig. 4A-B**). We next measured ATP-induced K^+^ current in macrophages expressing a dominant-negative Rab11a (Rab11a S25N). Cells treated with Rab11a S25N showed significant reductions in ATP-induced K^+^ current (**Fig 4C-D**), showing the requisite role for Rab11 a in inducing K^+^ current. Depletion of Rab11 a with siRNA in MDMs followed by assessment of NLRP3 showed that it decreased IL-1β release (Fig. 4E) and caspase 1 activation (**Fig. 4F-G**). Thus, Rab11a has a requisite role in mediating TWIK2 plasmalemma translocation and NLRP3 inflammasome activation.

To identify the *in vivo* role of macrophage-expressed Rab11a in regulating NLRP3 inflammasome activation and inflammation, we first depleted endogenous mouse lung macrophages with liposomal clodronate^41^, and subsequently carried out adoptive transfer (via i.t. route) of monocyte-derived macrophages with and without siRNA-mediated Rab11a depletion as illustrated in **Fig. 5A**. We then induced endotoxemia in recipient mice. The mice transplanted with Rab11a- depleted macrophages showed significantly reduced NLRP3 inflammasome activation (**Fig. 5B-D**) and inflammatory lung injury as assessed by neutrophil and macrophage infiltration in lungs (**Fig. 5E-F**) and quantification of myeloperoxidase (MPO) activity in lungs (**Fig. 5G**).

**Figure 5.**
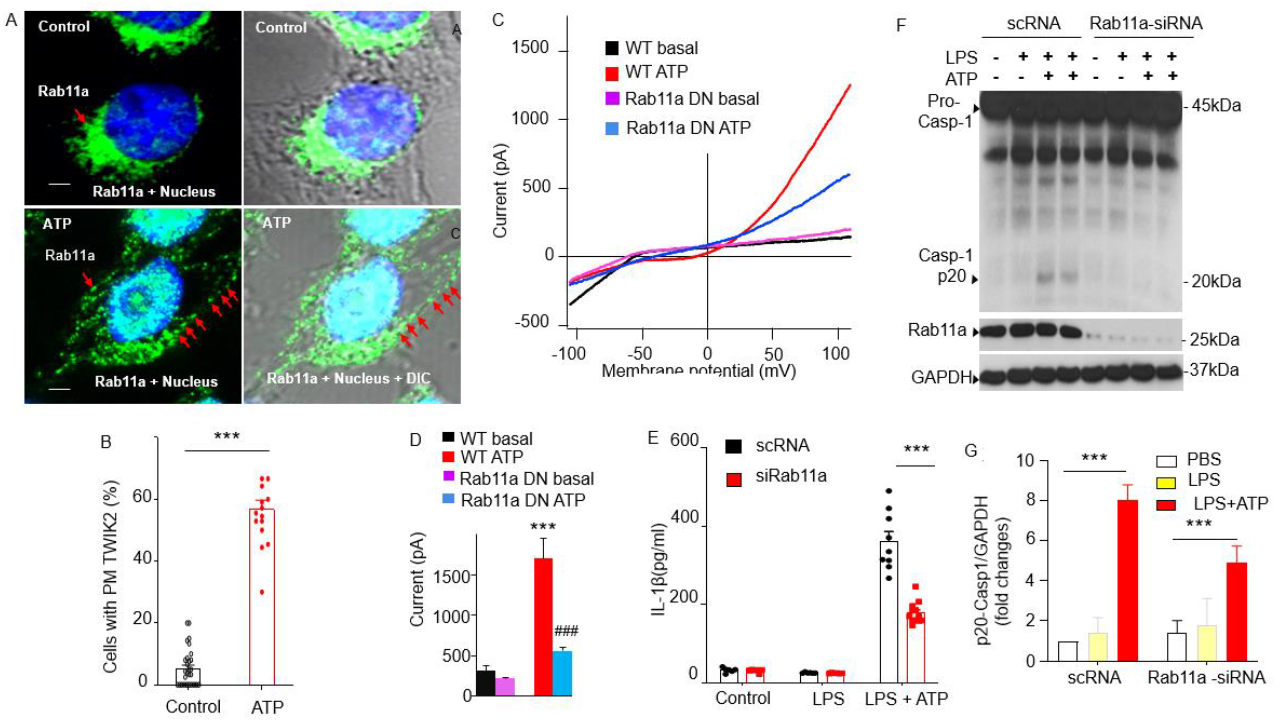
Rab11a mediates endosomal TWIK2 plasmalemma translocation and NLRP3 inflammasome activation on ATP challenge in macrophages. **A.** Confocal images of Rab11a immunostaining in mouse MDM before and after ATP challenge. Rab11a (green) distribution was identified with fluorescent immunostaining with Rab11a antibody (ab65200 from Abcam). Scale bar = 10μm. Note the dispersed distribution and plasmalemma translocation of Rab11a after ATP challenge (bottom panel). **B.** Summary of Rab11a plasmalemma translocation as shown in **A**. \*\*\**P*<0.001 compared with control group. **C-D**. Reduced ATP-induced K^+^ outward current in RAW 264 macrophages treated with dominant negative Ra11a (Rab11a DN) for 48 hr. Whole cell current was recorded with patch clamp as described in **Fig 3 C-D**. **C**. Representative I-V plots of whole cell current. **D**. Summary from experiments displayed in **C**. ***p < 0.001 compared with WT basal, n = 5). ###< 0.001 compared with WT ATP group, n = 5). Cells pretreated with Rab11a DN showed significantly decreased current induced by ATP. **E**-**G**. Inhibited NLRP3 inflammasome activation in MDMs treated with siRNA targeting mouse Rab11 a (siRab11 a). **E.** Reduced IL-1 β release in cells treated with siRab11 a. MDMs pretreated with siRab11 a for 48hr were primed with LPS (3 hr) and subsequently challenged with ATP (5 mM) for 30 min. IL-1β release in the supernatant was measured with ELISA. ***p < 0.001, n = 3. **F.** Representative Western blotting results from three independent experiments showing reduced Caspase 1 activation (reduced Casp-1 p20) and IL-1β maturation (reduced IL-1 β p17). MDMs pretreated with siRab11 a for 48hr were primed with LPS (3 hr) and subsequently challenged with ATP (5 mM) for 30 min. Cell lysates or pellets were immunoblotted with indicated antibodies (anti-TWIK2 or anti-IL1β). **G.** Quantification of results shown in **F**. ***p < 0.001, n = 3. Note the reductions Rab11a expression and Casp-1 p20 in cells treated with siRab11a. **Supplemental Figure 2**

**Figure 6.**
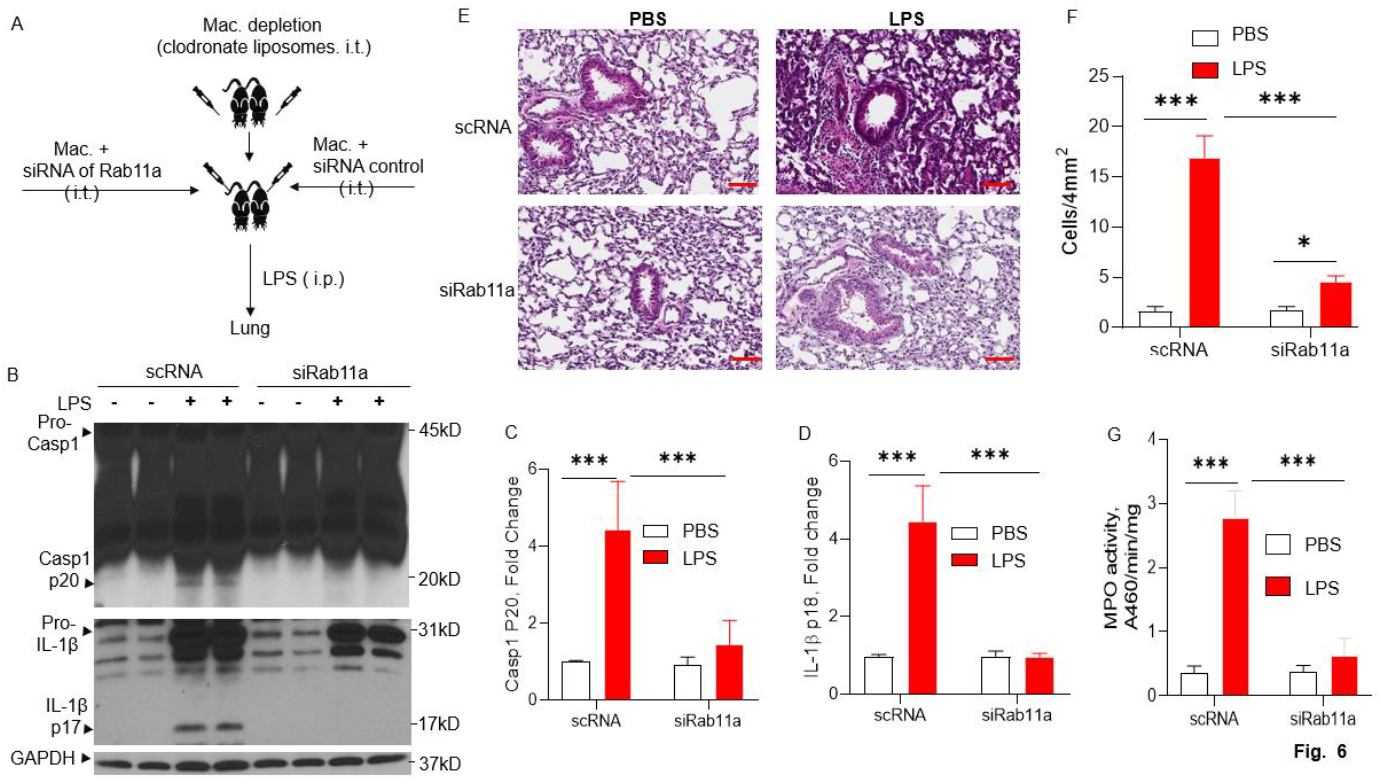
Rab11a deficiency in macrophages prevents sepsis-Induced NLRP3 inflammasome activation and inflammatory lung injury in mice. **A.** Schematic illustration of the experiments. Lung macrophages (Mac) were depleted with clodronate liposomes and then reconstituted via intratracheal route with MDMs treated with either siRNA of Rab11a or siRNA control as illustrated. The mice were injected with LPS (intra-peritoneal injection, i.p.) after 24 hr of macrophage reconstitution. Lungs were harvested for evaluation of NLRP3 inflammasome activation and lung inflammation. NLRP3 inflammasome activation (indicated by caspase 1 activation and IL-1β maturation) in the murine lung was assessed by immunoblotting (**B**) and quantified in (**C**) and (**D**) (***p< 0.001, n = 3). Representative H&E images of lung sections from three independent experiments are shown in (**E**, Scale bars: 200 μm.) and lung injury shown in **E** was evaluated by quantification of inflammatory cells in alveoli (per 4mm^2^ using the *Fiji* image analysis software) in **F** (*p < 0.05, ***p< 0.001, n = 3). Lung neutrophil infiltration was evaluated by MPO measurements of lung tissue shown in **G** (***p< 0.001, n = 3).

## DISCUSSION

The inflammasomes as the innate immune signaling receptors monitor the extracellular space and subcellular compartments for signs of infection, damage, and other cellular stressors^42^. NLRP3 inflammasome is a protein complex consisting of the inflammasome sensor molecule NLRP3, the adaptor protein ASC, and caspase 1^42,43^. NLRP3 formation is triggered by a range of substances generated during infection, tissue damage and metabolic imbalances^42,43^; however details of NLRP3 activation by events occurring at the plasma membrane remain unclear. NLRP3 activation is comprised of an initial priming phase involving NF-κB-dependent transcription of NLRP3 and pro- interleukin-lβ initiated by pro-inflammatory cytokines or via stimulation of Toll-like receptor (TLR) by agonists such as lipopolysaccharide (LPS)^44^. The second phase of NLRP3 activation is initiated by <u>P</u>athogen-<u>A</u>ssociated <u>M</u>olecular <u>P</u>atterns (PAMPs) or <u>D</u>anger-<u>A</u>ssociated <u>M</u>olecular <u>P</u>atterns (DAMPs) such as ATP, which ligates the purinergic receptor P2X7^45^ Based on studies on the structure and assembly mechanisms of NLRP3 complex, an endogenous, stimulus-responsive form of full-length mouse NLRP3 is a 12- to 16-mer double-ring cage held together by leucine-rich-repeat (LRR)-LRR interactions with the pyrin domains shielded within the assembly to avoid premature activation^9^. This NLRP3 form (double-ring cages of NLRP3) is predominantly membrane associated, such as endoplasmic reticulum - ER, mitochondria, Golgi Apparatus^10–12^ and the trans-Golgi network dispersed vesicles (dTGNvs), an early event observed for many NLRP3-activating stimuli^9^. Double-ring caged NLRP3 is recruited to the dispersed TGN (dTGN) through ionic bonding between its conserved polybasic region and negatively charged phosphatidylinositol-4-phosphate (PtdIns4P) on the dTGN. dTGNvs serve as a scaffold for NLRP3 aggregation into multiple puncta, leading to polymerization of the adaptor protein ASC, and thereby activating the downstream signaling cascade^13^ These results suggest a physiological NLRP3 oligomer on the membrane poised to sense diverse signals to induce inflammasome activation^9^ Evidence also shows that NLRP3 inflammasome is assembled and activated at the centrosome^14–17^, the major microtubule organizing center in mammalian cells, accounting for the singularity, size, and perinuclear location of activated inflammasomes^14^ However, little is known about the triggers at the plasma membrane initiating the downstream assembly and activation of NLRP3 complex.

Although it is known that a key upstream trigger of NLRP3 assembly and activation is potassium efflux^18–20^ via the potassium channel TWIK2^22^ which creates regional pockets of low potassium thus facilitating NLRP3 assembly, the nature of potassium efflux leading to activation of the NLRP3 complex is unknown. It was reported that cellular K^+^ efflux stabilized structural change in the inactive NLRP3, promoting an open conformation as a step preceding activation^23^. This conformational change appeared to facilitate the ensemble of NLRP3 into a seed structure for ASC oligomerization, a key step for NLRP3 inflammasome activation^23^. In the present study we examined how cells prevent continuous activation of the NLRP3 inflammasome despite TWIK2 being a continuously active background K^+^ channel^21^.

Electrophysiological characterization and understanding of the functional significance of TWIK channel family (TWIK1, TWIK2, and TWIK7) is impeded by the low or absent functional expression in heterologous expression systems^21^. TWIK1 was reported to be mainly located in intracellular compartments (such as pericentriolar recycling endosomes), and its transfer to the plasma membrane is tightly regulated^21^. A study also showed that TWIK2 generated background K^+^ currents in endolysosomes regulated the number and size of lysosomes in MDCK cells^46^. TWIK2 contains sequence signals responsible for the expression of TWIK2 in the Lamp1-positive lysosomal compartment^46^, and sequential inactivation of these trafficking motifs prevented the targeting of TWIK2 to lysosomes, thus promoting plasmalemmal relocation of the functional channel^46^. Here we determined the mechanisms of TWIK2 expression at the plasmalemma to address how the channel is functionalized in macrophages in the face of high intracellular K^+^ concentration. We showed that TWIK2 in macrophages was mostly distributed in endosomes but translocated on demand by ATP within 2 minutes to the plasmalemma by Rab11a mediated translocation. We observed that TWIK2 was primarily located in the endosomal compartment at rest, thus prevented TWIK2 mediated K^+^ efflux into the extracellular space to avoid unchecked NLRP3 activation. Upon ligation of the purinergic P2X7 receptor by extracellular ATP, however, Ca^2+^ influx via P2X7 activated the Ca^2+^ sensitive endosomal GTPase Rab11a to induce endosomal TWIK2 translocation to the plasma membrane. K^+^ efflux via plasma membrane translocated TWIK2 caused local K^+^ concentration ([K^+^]_in_) to decrease leading to NLRP3 inflammasome activation and phenotype transition of macrophages (**Supplemental Fig. 3**). Thus, endosomes served as “reservoirs” for the ion channel and their transport to plasmalemma upon stimulation.

There is precedence for this model. An increase in plasmalemma surface expression of G protein-activated inwardly rectifying K^+^ (GIRK) channels from recycling endosome functioned to modulate neuronal activity^24–26^ Recycling endosomes also served as intracellular storage compartments for the cardiac pacemaker channels - hyperpolarization-activated cyclic nucleotide-gated (HCN) ion channels HCN2 and HCN4 for rapid adaptation of their surface expression in response to extracellular stimuli^27^.

Endosomal membrane trafficking requires the coordination of multiple signaling events to control cargo sorting and processing and endosome maturation^47^. Several key regulators have been identified in endosome trafficking, such as the small GTPases (as regulators, initiating signaling cascades to regulate the direction and specificity of endosomal trafficking), Ca^2^, and phosphoinositides^47^. Here we found that the GTPase Rab11a was the most highly expressed in macrophages. Rab11a is thought to regulate the function of a special endosomal subpopulation, the recycling endosomes^39,40^. This heterogeneous tubular-vesicular compartment engaged in membrane trafficking, connects the endo- and exocytotic pathways^39,40^. Although Rab11 is prominent in recycling endosomes, other studies addressed its role in various intracellular domains, trans-Golgi network (TGN) and post-Golgi secretory vesicles^26,39,40^. Rab11a play a key role in mouse embryonic development through regulating the secretion of soluble matrix metalloproteinases (MMPs) required for cell migration, embryonic implantation, tissue morphogenesis, and innate immune responses^39^. Rab11a-null embryos formed normal blastocysts but died at peri-implantation stages^39^. We previously identified the role for Rab11a in regulating efferocytosis via the modulation of disintegrin and metalloproteinase (ADAM)17-mediated CD36 cell surface expression as a promising strategy for activating the resolution of inflammatory lung injury^48^. The present study shows a key role of Rab11a in mediating cycling endosomal TWIK2 plasmalemma translocation. Importantly as a test of functional relevance, adoptive transfer of Rab11a-deleted macrophages into mouse lungs after alveolar macrophage depletion prevented NLRP3 inflammasome activation and inflammatory lung injury. These results demonstrated that Rab11a in macrophages has a fundamental check-point role in TWIK2 plasmalemmal translocation and regulating NLRP3 inflammasome activation and endotoxemia-induced inflammatory lung injury.

## EXPERIMENTAL PROCEDURES

### Mice, cell cultures, and reagents

C57 black 6 (C57BL/6) mice were obtained from Charles River Laboratory. *Twik2^−/−^* mice was a generous gift from Dr. Lavannya M. Pandit (Baylor College of Medicine)^49,50^. *P2×7^−/−^* mice were purchased from Jackson Laboratory. All mice were housed in the University of Illinois Animal Care Facility in accordance with institutional and NIH guidelines. Veterinary care and animal experiments were approved by the University of Illinois Animal Care & Use Committee. For LPS-induced injury, mice received a single intraperitoneal dose (20 mg/kg) of LPS (*Escherichia coli* 0111:B4, L2630, Sigma). Mouse bone marrow monocyte derived macrophage (MDMs) were induced and cultured as described^51^. The mouse RAW 264.7 macrophage cell line was obtained from ATCC (TIB-71™) and was cultured and propagated as instructed by the manufacturer’s protocol. Caspase-1 antibody (p20, AG-20B-0042-C100) was purchased from AdipoGen Life Sciences. IL-1β antibody (AF-401-NA) was purchased form R&D systems. TWIK2-EGFP plasmid (pLV[Exp]-Puro-CMV>mKcnk6[NM_001033525.3]/3xGS/EGFP) was designed by and purchased from VectorBbuilder (VB200618-1166ypg). TWIK2 antibody (LS-C110195-100) was purchased from Life Span Bioscience. Antibody against Rab11a was purchased from Abcam (ab65200). Antibody against EEA1 (C45B10), LAMP1 (D2D11) and PDI (C81H6) were purchased from Cell Signaling Technology. LPS (E. coli 0111:B4, Ultrapure, tlrl-3pelps, used to treat cells) were obtained from Invitrogen. ATP-Na^+^ (A2383), Vacuolin-1(673000) and other chemicals were purchased from Sigma.

### Dynamic observations of intracellular TWIK2 plasmalemma translocation in macrophages challenged with ATP

Time-lapse video recording with confocal microscope was used to follow intracellular TWIK2 plasmalemma translocation in macrophages challenged with ATP. TWIK2-GFP plasmid were transfected into RAW 264.7 cells for 48hrs and cells were imaged with confocal microscope in the presence or absence of extracellular ATP. Video recording was initiated once cells were exposed to ATP (5mM) using a Zeiss LSM 710 confocal microscope using 488 nm lasers and a 63× objective lens (NA 1.3) and an emission bandwidth of 500 nm. Dynamic analysis of intracellular TWIK2 plasmalemma translocation was performed using Fiji software.

#### TWIK2 immunostaining in macrophages

RAW 264.7 macrophages were plated at a density of 300,000 cells on 25 mm coverslips. Following a 24h incubation, cells were treated with different concentrations of ATP for 30min, followed by five washes with PBS, and fixation for 20 min in 3% paraformaldehyde–PBS. Cells were blocked and permeabilized in 0.25% fish skin gelatin (Sigma), 0.01% saponin (Calbiochem, San Diego, CA) in PBS for 30min. Cells were stained with TWIK2 antibody for 1h, coverslips were washed, then incubated with goat antirabbit-AlexaFluor488 (Molecular Probes) for 1h, washed again and mounted in 4% n-propyl gallate, 25 mM Tris at pH 8.5 and 75% glycerol. Related localizations of TWIK2 were imaged with a Zeiss LSM 710 confocal microscope using 488 and 561 nm lasers and a 63× objective lens (NA 1.3) and an emission bandwidth of 500–535 nm. Images were acquired with LCS software and images were processed with Fiji software.

#### TWIK2 localization by immunoelectron microscopy

RAW 264.7 macrophages were collected and fixed by 4% paraformaldehyde and 0.15% glutaraldehyde in 0.1 M PB buffer for 1 h. Cells were subsequently washed with 0.1 M PB buffer, dehydrated with ethanol and embedded with L.R. White resin (Electron Microscopy Science, Hatfield, PA) in a vacuum oven at 45 °C for 48h. Sections (100 nm) were incubated with anti-TWIK2 primary antibody (LSBio, #LS-C110195-100) for 3 h and further incubated with 10-nm gold-conjugated secondary antibody, goat anti-rabbit IgG (H+L; Ted Pella Inc., Redding, CA) for 1 h. Cells were further stained with uranyl acetate and lead citrate and examined on a FEI Tecnai F30 at 300 KV.

#### Whole cell recordings

Electrophysiological recordings were obtained using a voltage-clamp technique. All experiments were conducted at room temperature (22 - 24°C) using an EPC-10 patch clamp amplifier (HEKA Electronik GmbH, Lambrecht, Germany) and using the Pulse V 8.8 acquisition program (HEKA Electronik GmbH, Lambrecht, Germany). Whole cell currents were elicited by using a ramp protocol with test pulse range from −110 to + 110 mV (200 ms in duration). The holding potential was 0 mV. The pipette solution contained (in mM): 120 K-glutamic acid, 2 Ca-Acetate Hydrate, 2 Mg-SO4, 33 KOH, 11 EGTA and 10 Hepes, pH 7.2. The bath solution contained (in mM): 140 Na-glutamic acid, 2 Ca-Acetate Hydrate, 1 Mg-SO4, 10 HEPES, pH 7.4. Whole cell capacitance was recorded as described^28^ Whole cell currents and capacitance were analyzed using IGOR software (WaveMetrics, Lake Oswego, OR).

#### Quantitative RT-PCR for fusion protein expression in macrophages

Total RNA of cultured MDMs was extracted using the RNeasy Micro Kit (Qiagen) according to the manufacturer’s instructions. RNA isolated from MDMs was converted to cDNA using the High-Capacity cDNA Reverse Transcription Kit (Applied Biosystems). Real-time PCR was performed using SYBR Green Master Mix on ViiAZ (Applied Biosystems) according to the manufacturer’s protocols. The following primers were used for PCR: Rab6a: forward, 5’-GATACTGCGGGTCAGGAACG-3’, and reverse, 5’- GCAGCAGAGTCACGGATGTAA-3’; Rab6b: forward, 5’-AACCCGCTGCGAAAATTCAAG-3’, and reverse, 5’-CGGTCTTCCAAGTACATGGTTT-3’; Rab11a: forward, 5’-AGGAGCGGTACAGGGCTATAA-3’, and reverse, 5’- ATGTGAGATGCTTAGCAATGTCA-3’; Rab11b: forward, 5’-GCTGCGGGATCATGCAGATAG-3’, and reverse, 5’-CACGGTCAGCGATTTGCTTC-3’; Rab27a: forward, 5’-GGCAGGAGAGGTTTCGTAGC-3’, and reverse, 5’-GCTCATTTGTCAGGTCGAACAG-3’; Rab27b: forward, 5’-TGGCTGAAAAATATGGCATACCA-3’, and reverse, 5’-CCAGAAGCGTTTCCACTGACT-3’; Synaptotagmin VII-1 (Syt7-1): forward, 5’-TTGGCTACAACTTCCAAGAGTCC-3’, and reverse, 5’- CGGGTTTAGATTCTTCCGCTTC-3’; Syt7-2: forward, 5’-CAGACGCCACACGATGAGTC-3’, and reverse, 5’-CTGGTAAGGGAGTTGACGAGG-3’; Vapm2: forward, 5’-GCTGGATGACCGTGCAGAT- 3’, and reverse, 5’-GATGGCGCAGATCACTCCC-3’; Vamp3: forward, 5’-CAGGTGCCTCGCAGTTTGAA-3’, and reverse, 5’-CCTATCGCCCACATCTTGCAG-3’. The data were analyzed using the comparative cycle-threshold (CT) method, where the amount of target is normalized to an endogenous reference gene, GAPDH.

#### Silencing Rab11a

Dominant-negative Rab11a (Rab11a S25N) was a gift from Dr. Guochang Hu (The University of Illinois at Chicago)^48^. The siRNA targeting mouse Rab11a (L-040863-01-0005) and a siRNA negative control were obtained from Horizon Discovery Ltd. Transient transfections of these dominant-negative Rab11 a and siRab11 a into mouse MΦs (RAW 264.7 cell line) were performed with Amaxa mouse macrophage nucleofector kit (VPA-1009, Lonza) and DharmaFECT 4 Transfection Reagent (T-2004-01, Dharmacon) according to the manufacturer’s protocol. To evaluate the efficiency of Rab11a silencing, Rab11a expression was examined with Western blot and inflammasome activation was examined by measuring p20 intensity via Western blot and IL-1β release through ELISA as mentioned above in cells 2-3 days after transfection.

#### NLRP3 inflammasome activation

Prior to experimental treatments, macrophages incubated with 1 μg/ml LPS as priming signal to induce NF-κB–dependent upregulation of pro-IL-1β and NLRP3 expression^52^. The cells were primed with LPS for 3 h at 37°C and then priming medium was replaced with normal culture medium. To evaluate the NLRP3 inflammasome activation, macrophages were stimulated with 5 mM ATP for 30 min at 37°C and then IL-1 β release in the medium or bath solutions were measured by ELISA and Caspase 1 activation was evaluated by Western blot using p20 antibody of Caspase 1. Briefly, cell-free supernatants were collected and then assayed for murine IL-1 β (MLB00C, R&D Systems) by ELISA kit (R&D Systems) according to the manufacturer’s protocol and the adherent macrophages were collected to generate whole-cell lysate. Cell lysate samples and lung protein samples from mice were subjected to SDS-PAGE and transferred to membrane for Western blot analysis using various primary antibodies (Caspase 1-p20 and IL-1β).

#### ELISA analysis

IL-1β in mice serum and cell-free supernatants were measured by Quantikine^®^ sandwich ELISA kits (R&D Systems) according to the manufacturer’s instructions. The cytokine concentration of the properly diluted or undiluted samples in 96-well plates was measured at 450 nm wavelength of absorbance, and calculated by GraphPad Prism linear regression analysis.

#### Myeloperoxidase (MPO assay)

The lungs were homogenized in 1 mL of PBS with 0.5% hexadecyltrimethylammonium bromide. The homogenates were sonicated, centrifuged at 40,000 × g for 20 min, and run through two freeze-thaw cycles. The samples were homogenized and centrifuged a second time. The supernatant was then collected and mixed 1/30 (vol/vol) with assay buffer (0.2 mg/mL o-dianisidine hydrochloride and 0.0005% H_2_O_2_). The change in absorbance was measured at 460 nm for 3 min, and MPO activity was calculated as the change in absorbance over time.

#### Alveolar macrophage depletion and reconstitution

Commercially available clodronate liposomes (Clodrosome) were administered directly into lungs of 10-week-old mice using a minimally invasive endotracheal instillation method. The mice were anesthetized by ketamine and xylazine (45 mg/kg and 8 mg/kg, respectively) and were suspended on a flat board and placed in a semi-recumbent position with the ventral surface and rostrum facing upwards. Using curved blade Kelly forceps, the tongue is gently and partially retracted rostrally and 50 μl of clodronate liposomes is placed in the back of the oral cavity, which is then aspirated by the animal. Control liposomes (50μl) alone were similarly administered in the control group. After 2 days of clodronate treatment, mice were reconstituted by i.t. instillation in a similar manner with differentiated MDMs (these MDMs have been treated with siRab11a for 48h) at dose of 2×10^6^ in a 50μl volume per mice. The mice were injected with i.p. LPS (20mg/Kg) after 24 h of macrophage reconstitution. The lungs were flushed and harvested after 24 h of LPS challenge.

#### Statistical analysis

Statistical comparisons were made using two-tailed Student’s t test for comparisons of two groups or one-way ANOVA followed by the Tukey’s post hoc pairwise multiple comparisons when appropriate with Prism 6 (GraphPad). Experimental values were reported as the means ± S.E.M (standard error of the mean). Significance between groups was determined using the t-test (two tails) and asterisks indicate a statistically significant difference with the number of experiments indicated in parentheses.

## ACKNOWLEDGMENTS

The work was supported by NIH grants P01-HL60678, P01-HL077806, T32-HL007829, R01-HL118068 and R01-HL90152.

## AUTHOR CONTRIBUTIONS

A.D., L.H., Y.K., and A.B.M. designed the studies; A.D., L.H., P.T. and B. Z. performed the experiments and the data analysis; A.D., L.H., Y.K., and A.B.M.. wrote the paper with critical input from the other authors.

---

**Supplemental Figure 1.**
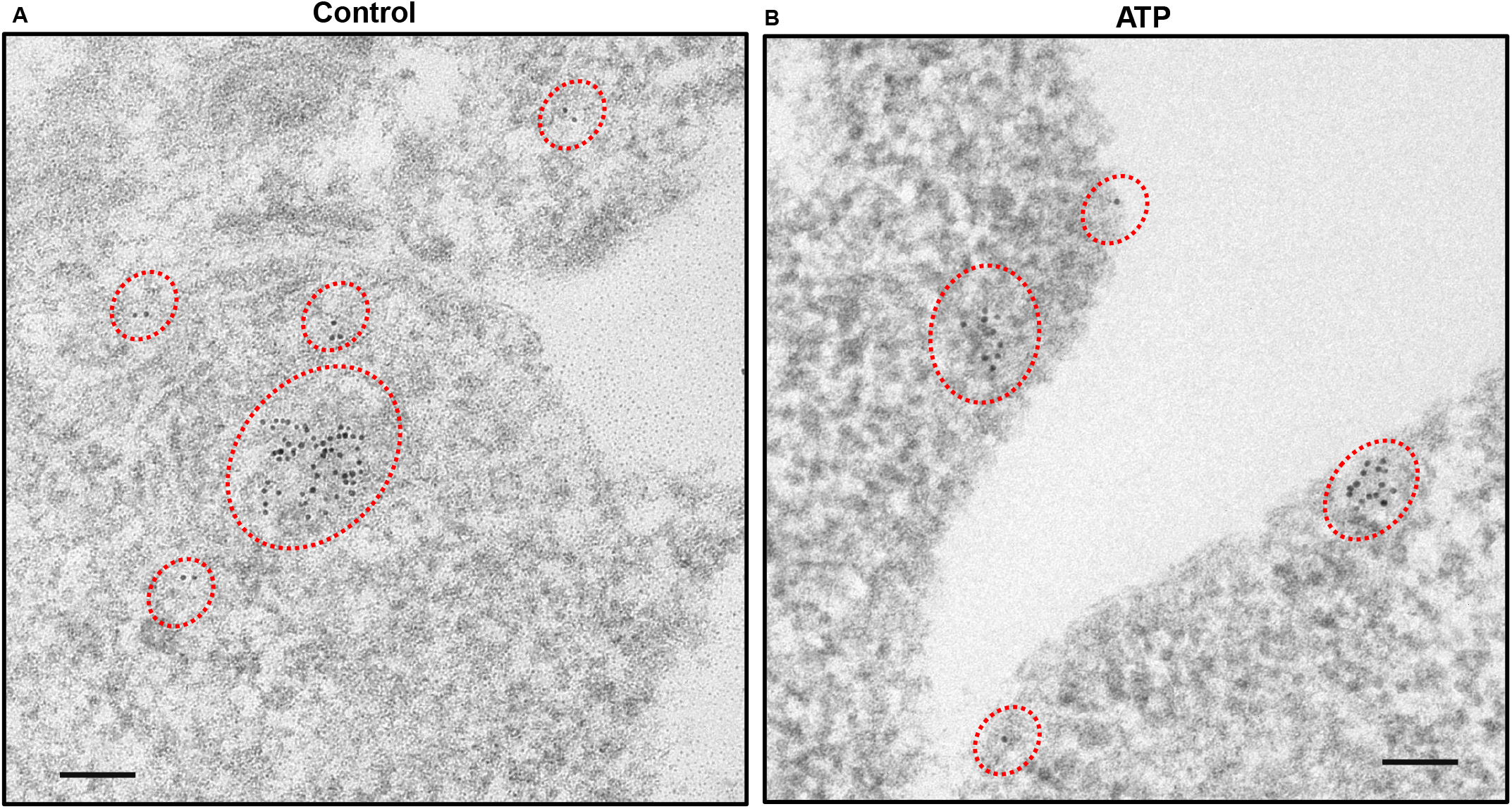
Electron microscopy assessment of TWIK2 plasmalemma translocation. TWIK2 plasmalemma translocation from immunogold labeling electron microscopy before (**A**) and after ATP (**B**) (5 mM, 30min) challenge in RAW 264.7 macrophages. TWIK2 (10 nm gold particles) was identified with anti-TWIK2 antibody as in **Fig. 1B**. Scale bar = 100nm. Note the vesicular structure outlined by the immunogolds in **A** and plasmalemma distribution of immunogold labeled TWIK2 after ATP challenge in **B**. Related to **Fig. 1C**.

**Supplemental Figure 2.**
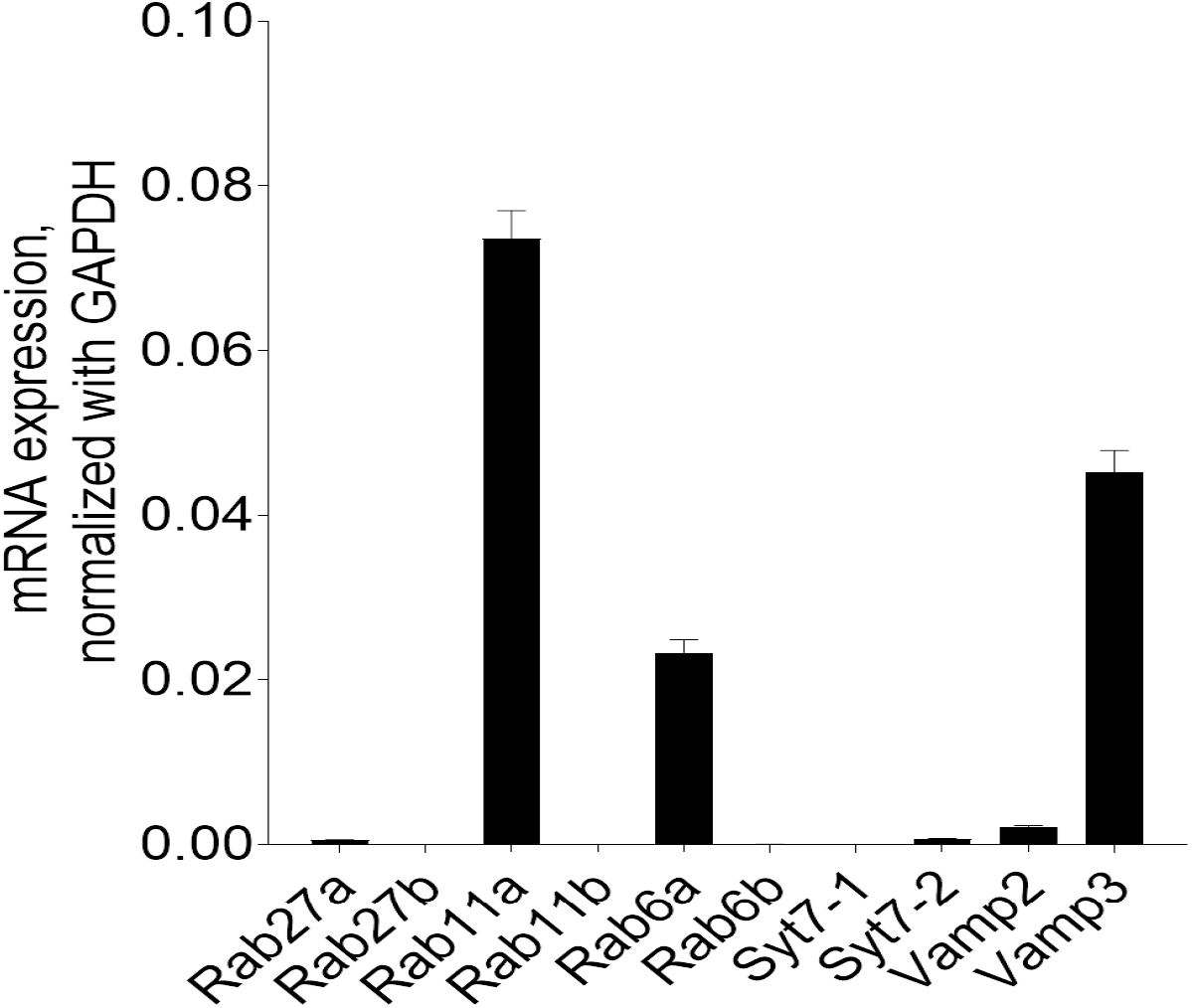
Relative mRNA expression of vesicle fusion proteins in MDMs. mRNA of various vesical fusion proteins were assessed by Qrt-PCR for the following: Rab27a and b; Rab11a and b; Rab6a and b; Synaptotagmin7-1 and 2(Syt7-1 and 2); Vamp2 and 3. The highest expression was seen in Ra11a (n = 3). Related to **Fig. 5**.

**Supplemental Figure 3:**
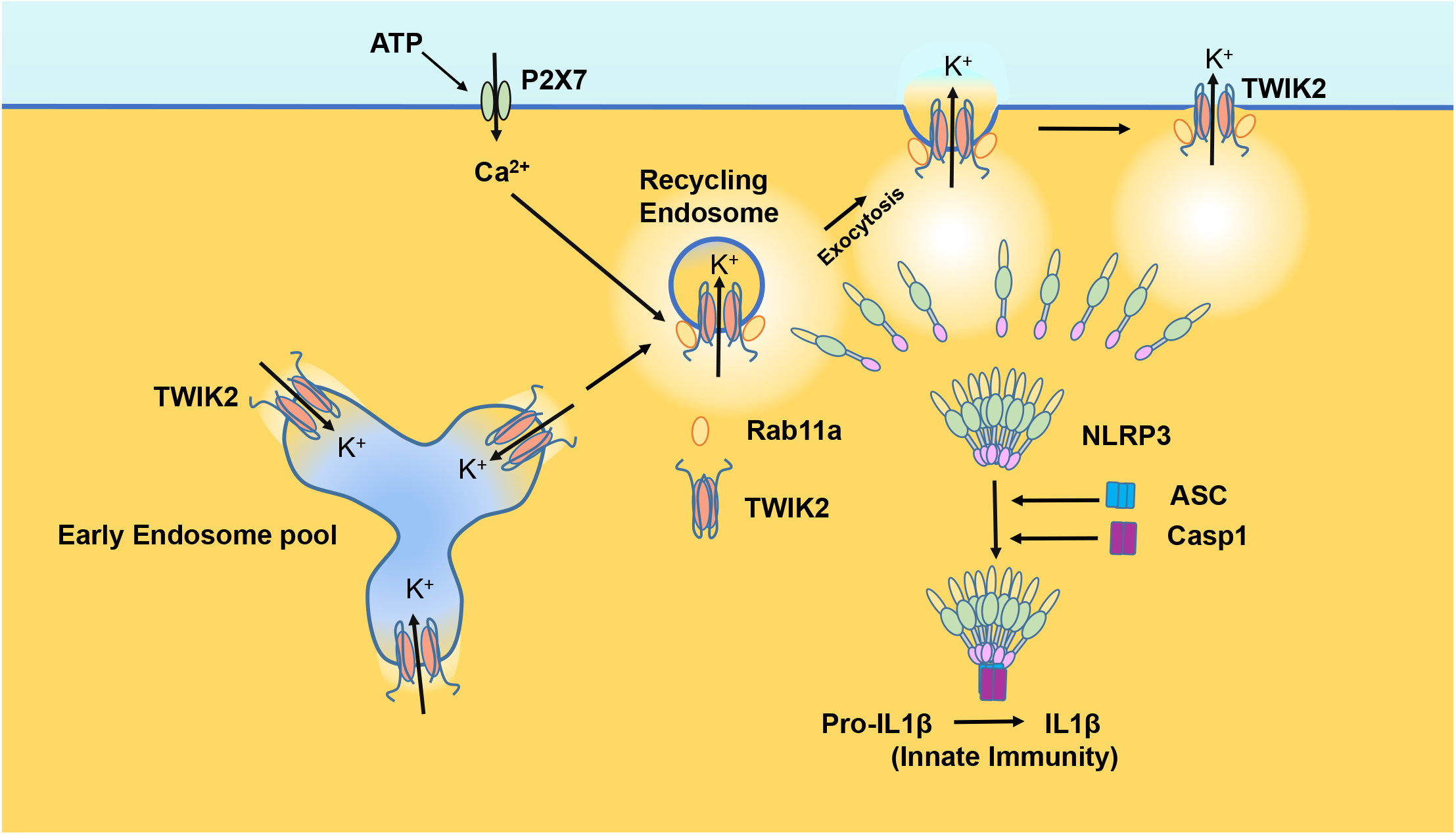
Endosomal TWIK2 plasmalemma translocation and resultant NLRP3 inflammasome activation. TWIK2 is basally active potassium channel in both early endosome (EE) and recycling endosome (RE). Extracellular ATP (e[ATP]) activates P2X7 and induces Ca^2+^ influx via P2X7 which activates Ca^2+^ sensitive Rab11a and causes recycling endosomal TWIK2 translocation to the plasma membrane. K^+^ efflux via plasma membrane translocated TWIK2 causes intracellular potassium concentration ([K^+^]in) to decrease leading to NLRP3 inflammasome activation. NLRP3 inflammasome activation thus leads to macrophage activation and promotes innate immunity. Related to **Fig 1** through **Fig 6**.

**Supplemental videos: Plasma membrane translocation of intracellular TWIK2 upon ATP challenge.** TWIK2-GFP plasmids were transfected into RAW 264.7 cells for 48hr and cells were imaged with confocal microscope in the presence (**video: TWIK2 GFP + ATP**) or absence (**vide: TWIK2 GFP - Control)** of extracellular ATP (5mM). Video recording was initiated once cells were exposed to ATP (5mM) using a Zeiss LSM 710 confocal microscope using 488 nm lasers and a 63× objective lens (NA 1.3) and an emission bandwidth of 500 nm. Dynamic analysis of intracellular TWIK2 plasmalemma translocation was performed using Fiji software. Related to **Fig. 1A**.

## Notes

### Competing Interest Statement

The authors have declared no competing interest.

